# Stabilising selection enriches the tails of complex traits with rare alleles of large effect

**DOI:** 10.1101/2024.09.12.612687

**Authors:** Anil P.S. Ori, Carla Giner-Delgado, Clive J. Hoggart, Paul F. O’Reilly

## Abstract

Establishing the relative contribution of common and rare variants to complex trait heritability is a key goal of biomedical research. Recent statistical genetics inference suggests that common variants explain most complex trait heritability, but little is known about how genetic architecture varies across the trait continuum. If rare variants make a small contribution to heritability but have their effects concentrated in the tails of complex traits, where disease typically manifests, then they may have a greater clinical impact than previously inferred. Here, we perform simulations using the forward-in-time simulator SLiM to generate population genetic and complex trait data, in which traits evolve under neutrality or stabilising selection and are entirely heritable. Recent studies suggest that stabilising selection is the dominant force shaping the genetic architecture of complex traits; this is consistent with our simulations here, since data simulated under stabilising selection more closely resembles real data. Moreover, we observe a shift of rare, large-effect alleles towards the tails of the distributions of traits simulated under stabilising selection. In our simulations, individuals in the tails of complex traits are, depending on the strength of selection, 10-20x more likely to harbour singleton or extremely rare alleles of large effect under stabilising selection than neutrality. Such an enrichment of rare, large-effect alleles in the tails of real complex traits subject to stabilising selection could have important implications for the design of studies to detect rare variants, for our understanding of the consequences of natural selection on complex traits, and for the prediction and prevention of complex disease.

---

Genome-wide association studies (GWAS) have demonstrated that most complex traits are highly polygenic^1,2^, with most of the contribution to heritability of this polygenicity coming from common, rather than rare (MAF < 1%), variants^3^. This presents a key challenge in precision medicine: namely, how to prevent and treat disease when its cause for most individuals appears to be the combination of hundreds or even thousands of risk alleles spread across genes of disparate function. It has been proposed that, despite polygenicity, there may be a relatively small number of ‘core genes’ most relevant to each disease^4,5^, or that different subgroups of patients may have risk clustered along specific genomic pathways that form one of multiple different disease aetiologies^6,7^.

While there has been intense focus on polygenicity in complex trait research^8,9^, there has been little investigation of whether levels of polygenicity vary across the trait continuum. However, polygenic risk scores based on large-effect rare variants have been shown to be more predictive of individuals in the tails of the population distribution^10^. If rare variants are highly enriched in complex trait tails, then there may be many individuals with a simple, monogenic trait or disease aetiology. This could have a profound impact on our approach to precision medicine. One factor that could affect how polygenicity varies across the trait continuum is the type and degree of natural selection that the trait is subject to. Under a scenario of neutrality, effect sizes should be independent of allele frequencies. However, under stabilising selection, considered the most common type of selection underlying complex traits^11,12^, trait variance in the population is reduced and large-effect alleles are restricted to low frequencies^13^. This could result in trait tails being enriched for rare alleles of large effect.

Here, we use the forward-in-time simulator SLiM^14^ to simulate population genetic data according to a Wright-Fisher (WF) model and parameters of mutation, recombination and effect size generation informed by empirical data. We perform simulations that produce complex traits based on polygenic, additive risk under neutrality and under varying degrees of stabilising selection. We benchmark the simulated genetic variation data with theoretical and empirical expectations and investigate how stabilising selection impacts variation in genetic architecture across the trait continuum, with a focus on the trait tails where disease typically occurs.

## Results

### Realistic data simulated under neutrality and stabilising selection

To investigate how the genetic architecture of a quantitative, polygenic trait is affected by stabilising selection, we implemented a forward-in-time simulation model, using SLiM^14^. Simulations were set-up to generate genomes corresponding to 10,000 individuals under a Wright-Fisher (WF) model^15^ of discrete, non-overlapping generations, within which mutations and recombination occur at genome-wide average rates and an individual’s phenotype value is calculated as the effect-size weighted sum of genome-wide alleles, with effect sizes drawn from a Gamma distribution (see Methods for details). To account for stochastic variation, we performed 100 replicates in each simulation scenario. Key parameters of the model used are shown in Fig. 1A and Fig. S1. During the burn-in period, which is performed under neutral evolution for 100k generation to reach an equilibrium, we observe an initial increase in genetic diversity (measured by heterozygosity) that plateaus over time (Fig. 1B), as expected. Stabilising selection is then implemented over the next 10k generations, during which genetic diversity reduces (Fig. 1B). On the phenotypic level, stabilising selection reduces the phenotypic population-variance (Fig. 1C), while the population-mean trait value stabilises once selection begins (Fig. 1D). The reduction in phenotypic variance occurs rapidly and drastically (Fig. 1E). Finally, stabilising selection prevents alleles of large effect from becoming common in the population (Fig. 1F), producing the typical “trumpet plot” pattern^16^ (blue points in Fig. 1F) that characterizes the relationship between allele frequency and effect size distribution observed in humans. Taken together, these observations are reflective of established features of genotype-phenotype data relating to traits under neutrality and stabilising selection, supporting the validity of our simulations and model implementation.

**Figure 1.**
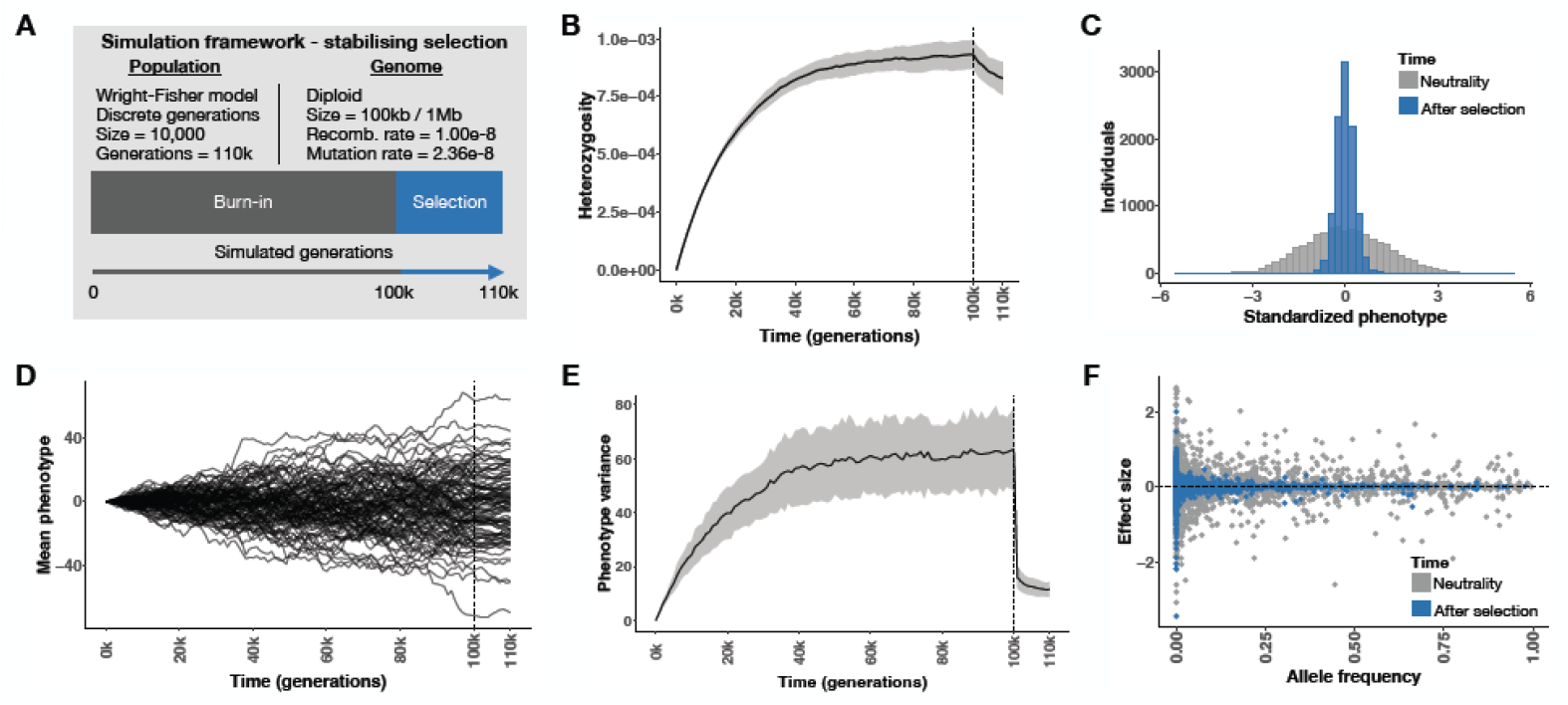
Basic characteristics of genetic variation data, simulated by SLiM, under neutrality and stabilising selection. (A) a schematic overview of model implementation and its key parameters. In panels B-F, results are shown in relation to genetic and phenotypic variation in the population before and after the period of stabilising selection (i.e. at generations 100k and 110k, respectively), based on 100 independent simulation replicates: (B) mean (black line) and standard deviation (SD – in grey) of population genome-wide heterozygosity over time (in generations), (C) the phenotype distribution in the population before and after selection from a single representative simulation, (D) the population-mean phenotype over time within each simulation replicate, (E) the population-variance phenotype over time (mean across replicates in black, SD in grey), (F) the relationship between variant allele frequency and effect size, before and after selection from a single representative simulation. The dotted vertical line in panels B, D and E denotes the start of stabilising selection at generation 100k.

### Stabilising selection increases the proportion of rare effect alleles

To further investigate the impact of stabilising selection on trait genetic architecture, we investigated the site frequency spectrum (SFS) across trait-associated variants, before and after selection. Comparing changes in the SFS between the start and end of stabilising selection, we observe a significant increase in the proportion of rare variants (MAF < 1%) – each of which has an effect in these simulations – by the end stabilising selection (Fig. 2A). At the end of the burn-in period of 100k generations under neutrality, 56.7% (SD=3.4%) of variants (across replicates) were rare, while this had increased to a mean of 64.7% (SD=5.2%) after 10k generations of stabilising selection. On average, taking into account the reduction in total number of variants under stabilising selection, rare variants were increased 1.41x (95% CI: 1.35-1.49, P < 2.2e-16) after selection compared to neutrality. The observed effects were more pronounced when considering extremely rare variants, with MAF < 0.05% (i.e. count ≤10 here). Higher resolution longitudinal analyses show that rare variants are relatively enriched in the population at the start of the burn-in period (Fig. 2B), when new mutations enter the population, reducing as equilibrium is reached. During stabilising selection, rare variant enrichment gradually increases and plateaus at the end of selection (Fig. 2B).

**Figure 2.**
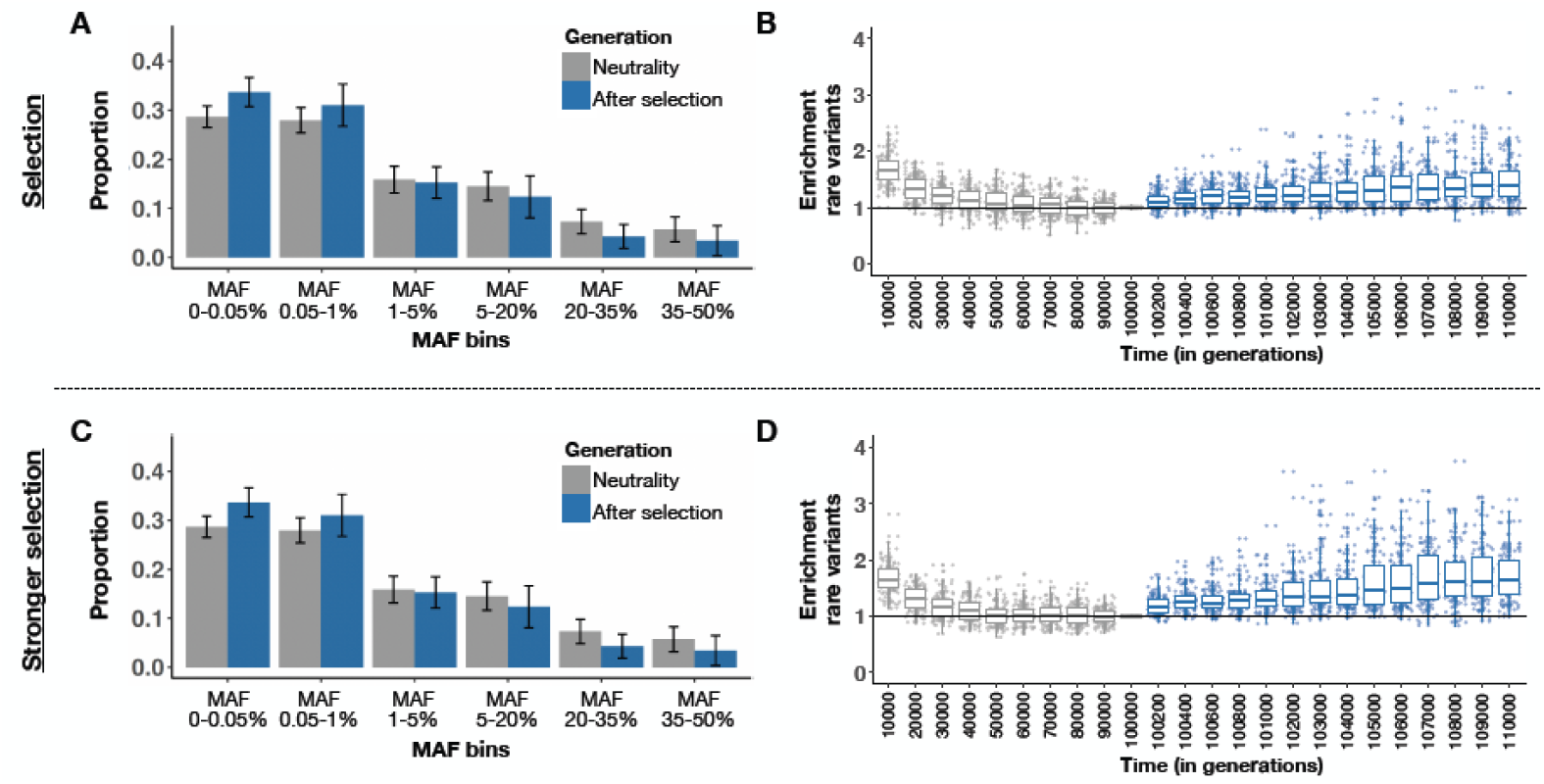
Changes in the site frequency spectrum (SFS) of causal variants due to stabilising selection. The top panels (A & B) illustrate results from simulations with selection (1x strength), while the bottom panels (C & D) show results with stronger selection (100x strength). Panels A and C show the mean (bars show standard errors) proportion of variants in minor allele frequency bins across 100 simulation replicates before (generation 100k - in grey) and at the end of selection (generation 110k - in blue). The MAF threshold of 0.05% includes extremely rare variants with a count of 10 (10/20,000 genomes = 0.05%) or fewer in the population. Panels B and D show the enrichment of rare variants (defined as MAF < 1%) over time. Enrichment is calculated as an ‘odds ratio’ defined by the rare:common ratio at a given generation divided by that ratio at the start of selection (at generation 100k). Box plots are shown summarising the distribution of enrichment across simulation replicates at a given generation (grey = before selection, blue = during selection).

In separate simulations performed under stronger selection (see Methods), we observe a similar trend with even stronger enrichments of rare variants. The fraction of rare variants again gradually increased during stabilising selection, reaching 68.9% (SD=5.5%) by the end (Fig. 2C & 2D). This represents a rare variant enrichment of 1.68x (95% CI: 1.60-1.77, P < 2.2e-16) compared to neutrality, which is significantly greater than under regular stabilising selection (P=2.23e-6). These observations indicate that the proportion of rare effect alleles increases when the corresponding trait is subject to stabilising selection, to a degree dependent on the strength of selection.

### Stabilising selection enriches trait tails with large-effect rare alleles

Our simulations have shown that stabilising selection limits alleles of relatively large effect to lower allele frequencies (Fig. 1F) and shifts the site frequency spectrum of effect alleles towards rarer frequencies (Fig. 2). Since large-effect alleles are more likely to ‘shift’ individuals towards, or into, the trait tails, and are more likely to be rare under stabilising selection, then we hypothesise that individuals in trait tails are more likely to carry rare alleles of large effect that are yet to be removed by selection. To evaluate our hypothesis, we calculate the enrichment of different classes of rare alleles, stratified by MAF and effect size, carried by individuals across the trait continuum under selection compared to neutrality. For the purpose of testing and throughout the rest of this manuscript, we define the tails as the lower and upper 1% of the trait distribution, which, while arbitrary, approximates the prevalence of many common diseases. Observed patterns are expected to be the same in both tails given the symmetry of effect sizes and selection coefficients simulated, but we report results separately here since these should be more reflective of real data results that focus on specific tails.

After stabilising selection, individuals carry 1.93x (95% CI: 1.68-2.27, P=4.2e-15) and 1.97x (95% CI: 1.70-2.27, P=5.1e-15) more rare alleles (MAF < 1%) than under neutrality in the lower and upper tail, respectively (Fig. 3A). We observe more pronounced enrichment in individuals carrying large-effect rare alleles, defined as variants with an effect size in the top decile of effect sizes. Individuals at the upper tail, for example, carry 4.29x (95% CI: 3.34-5.52, P=5.3e-20) more large-effect rare alleles after selection compared to neutrality, with similar enrichment observed in the lower tail. Furthermore, we observe that large-effect rare alleles are depleted in individuals at the center of the trait distribution after selection (OR=0.54, 95% CI: 0.43-0.67, P=1.5e-07).

**Figure 3.**
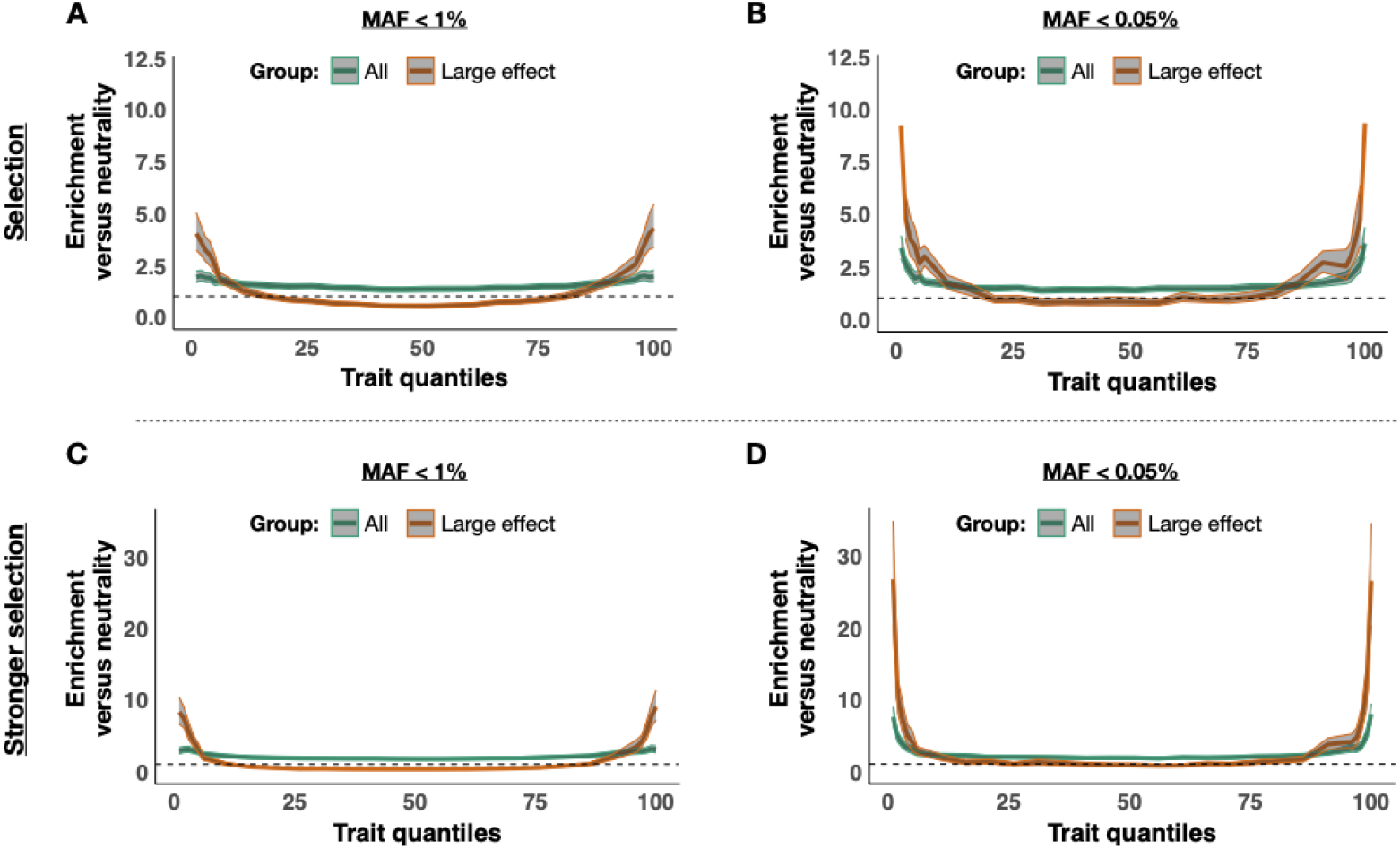
Enrichment of rare alleles across the trait continuum under stabilising selection. The enrichment of rare alleles after selection (at generation 110k) is shown across quantiles of the phenotype distribution for simulations with selection (upper panels) and stronger selection (lower panels). Panels A and C show results for rare variants with MAF < 1%, while panels B and D shows results for extremely rare variants with MAF < 0.05% (allele count ≤ 10). Enrichment (Y-axis) is calculated, in each trait quantile, as an odds ratio defined by the ratio of rare:total alleles carried by individuals after selection divided by that ratio under neutrality (i.e. before selection at generation 100k). For each panel, the enrichment is shown across the trait distribution with corresponding 95% confidence intervals. In green are results for all variants at each MAF threshold, while orange shows results for alleles with large effect (defined as alleles with effects in the top decile of absolute effect size). The dashed horizontal lines represent an odds ratio of one (i.e. no difference between neutrality and selection).

We observe an even greater enrichment of extremely rare alleles (MAF < 0.05%) in the trait tails compared to those that are moderately rare (MAF < 1%). Individuals in the upper tail have an enrichment of extremely rare alleles of 3.61x (95% CI: 2.97-4.39, P=3.5e-23) under selection compared to neutrality (Fig. 3B), while the enrichment of those that are both extremely rare and have a large effect is 9.33x (95% CI: 7.01-12.42, P=1.3e-23).

For all groups of rare variants, simulations with stronger selection produced even more pronounced enrichments (Fig. 3C & 3D). Extremely rare alleles are, for example, enriched 9.03x (95% CI: 7.16-11.40, P=2.0e-34) in the upper tail, and those of large effect are enriched 26.59x (95% CI: 20.30-34-81, P=3.1e-35). Similar qualitative patterns of rare-variant enrichment in the tails were observed in simulations in which we implemented different genetic effect-size generating distributions, with variation in the degree of enrichment between distributions. Gamma distributions with a smaller mean produced higher enrichment in the tails, while a Gaussian distribution produced distinctly lower enrichment (Fig. S2-S5). Together, these findings suggest that the contribution of rare variants to tail genetic architecture increases under stabilising selection, and that the increased contribution is most pronounced for extremely rare variants of large effect.

### Stabilising selection reduces PRS signal in trait tails

Our simulations have shown that stabilising selection can affect how genetic architecture varies across the trait continuum, in particular producing an enrichment of large-effect rare alleles in the tails. Next, we investigate the potential impact of this on the polygenic risk score (PRS) corresponding to the trait. First, PRS are computed for each simulated ‘individual’ as the effect-size weighted sum of their effect (i.e. derived) alleles genome-wide, in which effect sizes are known from the simulation and all common SNPs (MAF > 1%) are included (see Methods). Next, we investigate how the PRS varies across the trait continuum under neutrality (Fig.4A) and stabilising selection (Fig.4C). We also calculate ΔPRS – the difference between the observed and expected PRS – in the tails of the trait distribution (see Methods and Fig. S6), since an enrichment of rare alleles is associated with a reduction in common alleles, by definition, which should result in reduced (absolute) PRS relative to expectations. Under neutrality, we observe an average ΔPRS of 0.07 (SD=0.16; *P* = 2.5×10^5^) and 0.12 (SD=0.42; *P* = 6.0×10^-3^) in the lower (bottom 1%) and upper (top 1%) tail, respectively, across 100 simulation replicates (Fig. 4B). Under stabilising selection, we observe an average ΔPRS of 1.24 (SD=1.14; *P* < 2.2×10^-16^) and 1.39 (SD=1.09; *P* < 2.2×10^-16^) in the lower and upper tail, respectively (Fig. 4D). This indicates substantial “regression-to-the-mean” of PRS in the tails of the trait distribution due to enrichment of rare alleles of large effect, which contribute to extreme trait values but are not included in standard PRS based on common variants. In simulations with stronger selection (100x), we observe an even greater regression-to-the-mean tail effect of 2.74 (SD=1.30; *P* < 2.2×10^-16^) and 2.66 (SD=1.29; *P* < 2.2×10^-16^) in the lower and upper tail, respectively. In simulations that implement different genetic effect-size distributions, we observe similar patterns of regression-to-the-mean with a Gaussian distribution but, again, with a smaller impact compared to when effect sizes follow a Gamma distribution (Fig. S7, Fig. S8).

**Figure 4.**
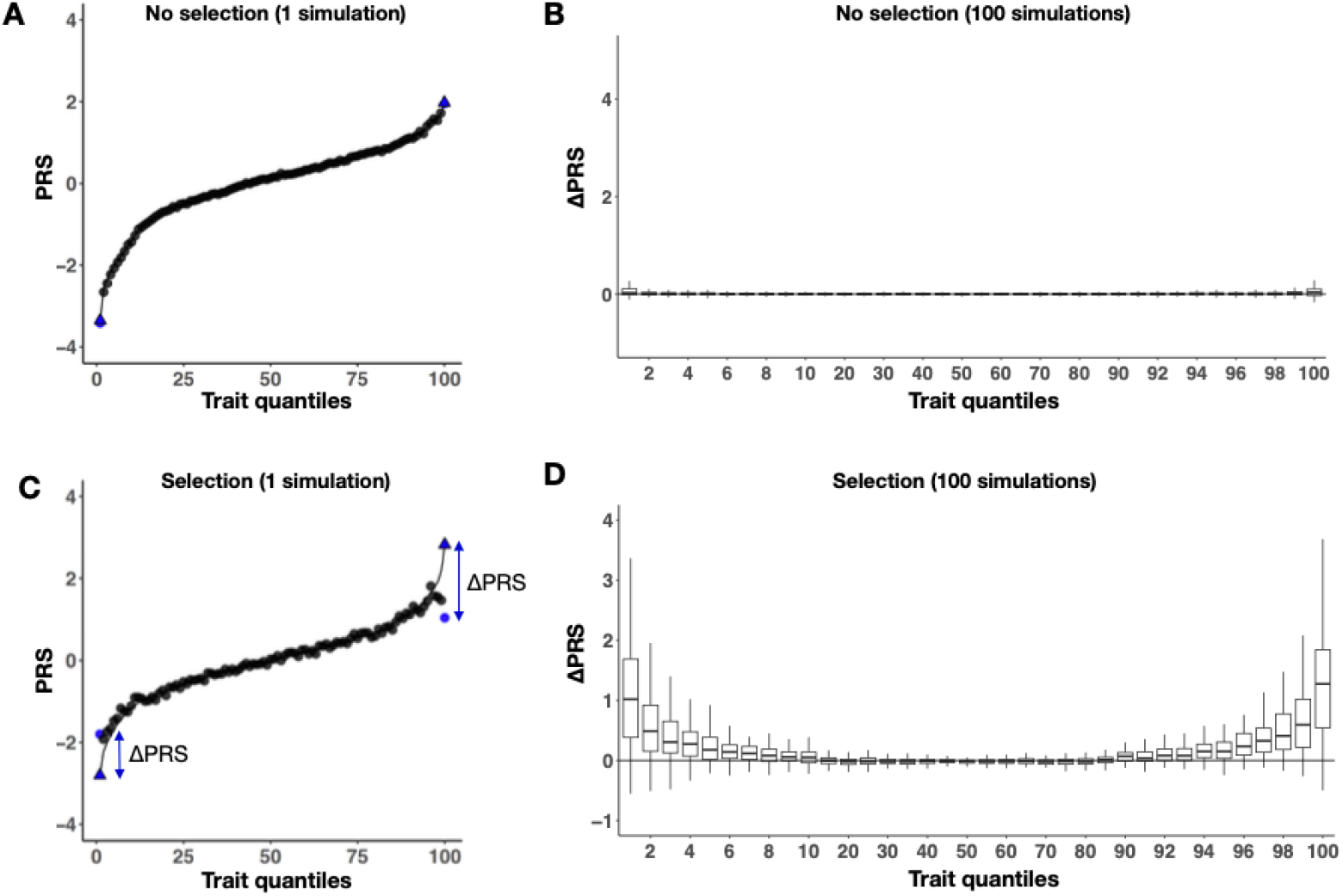
Variation in PRS across the trait continuum under neutrality and stabilising selection. The average PRS (see Methods) of individuals in each percentile of the trait distribution for a single representative simulation (A under neutrality; C under stabilising selection) and for 100 simulation replicates summarised as box plots (B under neutrality, D under stabilising selection). In panels A and C, the curved line shows the expected PRS based on a regression of PRS on trait between the 10^th^ and 90^th^ percentiles of the trait distribution (see Methods), while the tail percentiles (1 and 100) are coloured blue for emphasis, with the observed value shown as a disc and expected value as a triangle. ΔPRS = |PRS_exp_| - |PRS_obs_|.

### PRS signal is mostly recovered by inclusion of extremely rare variants

To further characterise the PRS regression-to-the-mean effect observed in the tails under stabilising selection (Fig.4), we investigated the extent to which it can be explained by different classes of rare variants. We recalculated each simulated individual’s PRS with the addition of rare variants within specific MAF and effect size bins and assessed the change in ΔPRS. We refer to these PRS augmented with rare variants as “rescue models”, given the expectation that they should contribute to recovering the expected PRS signal. We defined rare variants at three MAF thresholds: singletons, variants with MAF < 0.05% (count ≤ 10), and all variants with MAF < 1%. In addition, we stratify these three groups by two effect size thresholds: any effect size and only large effect sizes (defined by variants in the top decile based on absolute effect size). We find that, in our simulated data, a majority of ΔPRS can be explained by relatively few large-effect variants that are extremely rare in the population (Fig. 5). Under stabilising selection, variants with a large-effect allele of count ≤ 10 explain, on average, just over half of ΔPRS, with the rare alleles carried by approximately 14% of individuals in the tails of the trait distribution. In simulations with stronger selection, we observe that these variants can explain up to two-thirds of ΔPRS in the tails, with approximately 28% of individuals being a carrier. In comparison, under neutrality, only 1-2% of individuals in the trait tails carry such variants, indicating an enrichment of 10-20x depending on the strength of selection.

**Figure 5.**
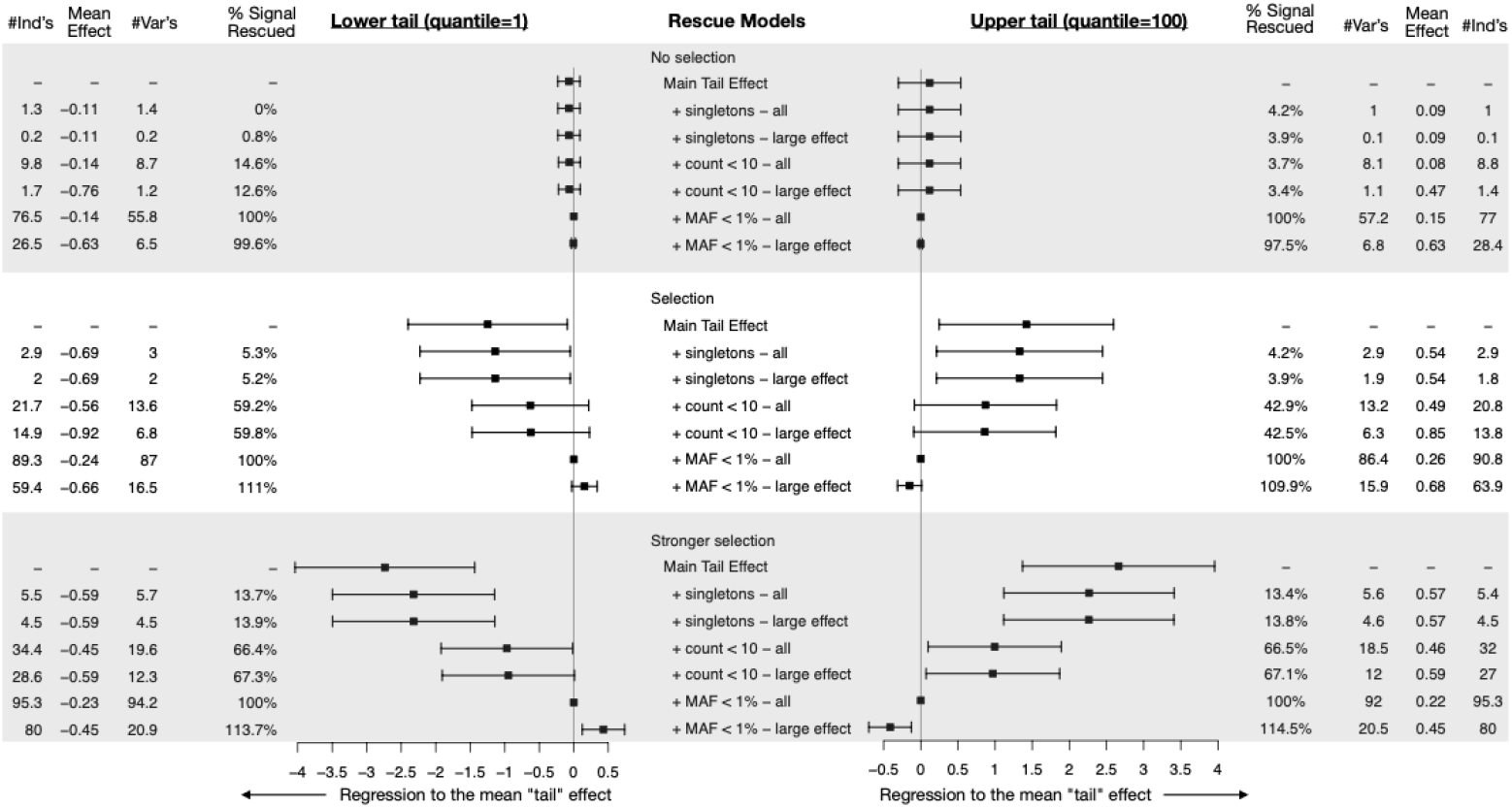
The contribution of rare variants of different classes to ΔPRS. Results show ΔPRS rescue experiments across simulation models under neutrality (top panel), selection (middle panel), and stronger selection (bottom panel). PRS was recalculated by including (large effect) singletons, alleles with a count ≤ 10 (MAF ≤ 0.05%), and variants with MAF < 1%. ΔPRS was also recalculated to determine the change in regression-to-the-mean tail effect and the corresponding % rescued. Results are shown in forest plots with squares denoting the mean and whiskers the SD of the regression-to-the mean across 100 simulations. Forest plots on the left show the distribution of ΔPRS at the lower trait tail (quantile 1) for each rescue model, while forest plots on the right show the results for the upper tail (quantile 100). Left and right columns show the average proportion of main regression to the mean effect that is rescued (% Signal Rescued), the average number of variants (#Var’s) within each tail (i.e. within the 100 individuals in that quantile), the average effect size of these variants (Mean Effect), and the average number of individuals (#Ind’s) that carry at least one of the effect alleles from these variants. Averages are calculated across 100 simulation replicates.

Singletons of large effect explain an estimated 4-5% of ΔPRS under stabilising selection and approximately 14% under strong stabilising selection. In the latter case, this corresponds to 4 or 5 individuals in each tail (comprising 100 individuals) with a singleton that contributes significantly to the observed ΔPRS.

As a positive control, we confirmed that the inclusion of all rare variants (MAF < 1%) rescued 100% of the tail effect in all simulations and that this appeared to be essentially due to rare variants of large effect. In fact, we note that when considering all rare variants of large effect under selection, more than 100% of the tail effect is rescued: this is because a depletion of rare variants of large-effect in the centre of the distribution calibrates the expected PRS to be lower than the true full PRS, since the expectation is based on the PRS in the centre (10%-90%) of the trait distribution.

Our rescue experiments demonstrate that, as a consequence of stabilising selection, extremely rare, large-effect variants are a key component of the genetic architecture of complex trait tails according to our simulations. By augmenting PRS with the inclusion of a handful of extremely rare, large-effect variants, a substantial fraction of the genetic risk of individuals in the tails can be captured.

## Discussion

Understanding the genetic architecture of human traits is essential for gene discovery and the utilisation of genetic information for disease prevention and management in clinical care^17^. Recent studies have pointed to the key role of stabilising selection in shaping complex trait genetic architecture^11,12,18,19^. Using forward-in-time simulations, we demonstrate that stabilising selection can have a substantial impact on the genetic architecture of a polygenic quantitative trait, particularly in the tails of the trait distribution. We find that, while stabilising selection reduces trait variation in the population, it increases the proportion of variants that are rare compared to that under neutral evolution. Importantly, this enrichment is greatest in the tails of the trait distribution and is particularly pronounced for extremely rare, large-effect variants. This suggests that, under stabilising selection, a significant proportion of trait variance in the tails can be explained by a relatively small number of genetic variants. This implies that the trait tails effectively have reduced polygenicity, both relative to that under neutrality and compared to the rest of the trait distribution.

Our work highlights a previously unreported, but potentially critical, consequence of stabilising selection. While it is known that stabilising selection reduces genetic variation in the population, limiting most large-effect variants to low frequency, we find here that it can also produce an enrichment of large-effect, rare variants specifically in the trait tails. This occurs because trait variation significantly reduces under stabilising selection, which causes individuals that carry large-effect variants to have a greater probability of being in the trait tails. The effect alleles of these variants arise newly in the population and, most likely, segregate in the population for a limited number of generations, having a disproportionate impact on extreme trait values, before being removed from the population due to negative selection pressure in the tails caused by the trait being subject to stabilising selection^13^. Accounting for such evolutionary processes, and their consequences for genetic architecture, will be imperative for optimal study design of risk variant detection and genetic risk prediction. As sample sizes of genome-wide association studies (GWAS) grow, strategies that jointly model effect size and allele frequency distributions, and that explicitly incorporate parameters of selection into PRS calculation^20,21^, should yield greater predictive performance. Including putative rare pathogenic variants in PRS models have shown promise in identifying individuals at risk for disease^22,10^ and our results suggest that this approach could be particularly powerful for traits that have been subject to stabilising selection. Finally, models that calibrate an individual’s genetic risk based on the expected variability in genetic architecture across the trait distribution may help to improve predictive performance further.

Our study includes several limitations. While we have implemented a sophisticated simulation model that leverages an established population genetic framework, we make multiple simplifying assumptions about population dynamics, genomic features, and trait characteristics. Firstly, we assume constant mutation and recombination rates, which are known to vary locally substantially in humans, although we do not expect this to have a qualitative impact on results here. Secondly, in our primary analysis, we assume that genetic effect sizes are drawn from a Gamma distribution because this reflects the heavy-tailed distributions typically modelled in the field^23,24^ and establishing true effect-size generating distributions is highly challenging. In our sensitivity analyses, we observed that genetic effect sizes drawn from gamma distributions with smaller means produce greater enrichment of rare variants in the trait tails after stabilising selection, while a Gaussian distribution produced relatively weak enrichment in the tails and correspondingly small regression-to-the-mean of the PRS signal. Further work should expand the space of effect-size distributions considered, although we expect the gamma distributions assumed in the primary analyses to reflect real genetic effects sufficiently well to have produced qualitatively realistic patterns. Thirdly, we simulate a genome of size 100kb for computational efficiency and do not investigate the impact of varying genome size. Target genome size could be an important factor in observed patterns of genetic architecture^25^ and so should be explored in follow-up work. Fourthly, we only model a single quantitative trait under a polygenic, additive model and did not explore the role of non-genetic or non-additive factors, nor did we investigate pleiotropic effects, known to be widespread for human complex traits^26,25^. Fifthly, we only explore the effects of stabilising selection, and while this is considered a key driving force shaping complex traits^11,12^, future work should consider other forms of selection, such as directional selection. Despite these multiple simplifying assumptions, our simulations show the patterns of genetic variation that can emerge across polygenic trait distributions when subject to selection, which can be extended and investigated further in real-world data^27^.

In summary, we demonstrate via simulation how the genetic architecture of a trait can vary substantially across the trait continuum when the trait is subjected to stabilising selection over a prolonged period. We find that rare variants, especially those of large effect, can be particularly enriched in trait tails after selection and contribute to a dramatic reduction in average absolute PRS values of individuals in the trait tails. Much of the observed reduction in PRS can be explained by extremely rare variants, suggesting that many individuals have extreme trait values due to a single or a few rare alleles of very large effect. Our findings have implications for gene mapping study design and for the genetic prediction of human traits, particularly of diseases, which typically manifest in the tails of trait distributions.

## Methods

### Wright-Fisher model of stabilising selection

We performed forward-in-time genetic simulations using a Wright-Fisher (WF) model^15^ as implemented in SLiM v4.0^14^. Simulations were initialized with a genomic sequence of 100kb in with a uniform recombination rate of 1.00 × 10^-8^ cross-over events per base pair (bp) per generation and a mutation rate of 2.36 ×10^-8^ per bp per generation. Causal effects sizes of new mutations were sampled from a gamma distribution in our primary analyses. A gamma distribution has been used in other models of human evolution^23^, reflecting a long-tail distribution that combines mostly small effects with a small number of relatively large effects. The gamma was parameterized with a shape of 0.186, which was extrapolated from the best shape estimate of the distribution of fitness effects for nonsynonymous mutations based on 1000 Genomes data.^24^. The mean of the gamma distribution was parameterized using three different values (i.e. −0.05, −0.10, −0.20) to reflect a range of effect sizes close to that observed in genetic association analyses of common diseases. Each mutation drawn was multiplied by δ, with δ∈{-1,1} with equal probability, to reflect a symmetrical effect size distribution commonly seen with complex traits. For our main simulation analysis, the effect-size generating distribution was defined as a gamma with mean=−0.10 and shape = 0.186. Sensitivity analyses were conducted using the two other gamma distributions, as well as using a Gaussian distribution parameterized as N(0,1) to assess how results vary across different genetic effect-size generating distributions.

Trait heritability was set to 1.0 with all variants contributing to the trait. At each generation, an individual (*i)* their phenotype (*y*) is defined by the sum of the number of alleles (*X*) of each mutation (*j*), weighted by the effect size of the allele (*β*);

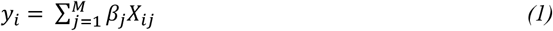

where M is the total number of mutations. We simulated a WF model with a diploid population of 10,000 individuals (N) that remained constant with discrete, non-overlapping generations. To reach an equilibrium state of mutation–drift balance, we first initialized a burn-in period of 10N generations with random mating. After neutral genetic diversity was achieved, stabilizing selection was introduced for 1N generations through a Gaussian fitness function,

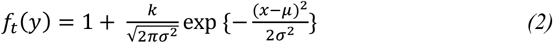

where *y* is the phenotype of the individual at generation *t, μ* is the phenotypic mean of the population at *t=10N* and *σ* the phenotypic standard deviation at the start of selection (i.e., *t*=10N). The factor *k* (i.e. to increase strength of selection) parameter was set at 1. To implement stronger selection, we used *k*=100. The fitness of each individual was calculated and updated at the end of each generation. Stabilizing selection was simulated until generation 11N after which the simulation was completed, unless otherwise mentioned. We ran 100 independent simulations and extracted information about the population every 1000 generations, which included the phenotypic mean and standard deviation of the population, average per bp heterozygosity, statistics of mutations present in the populations, including allele frequencies and effect sizes, and the mutational profile of each individual.

### Quantification of rare variant contribution to trait genetic architecture

To investigate trait genetic architecture, we first measured changes across the site frequency spectrum (SFS) before and after stabilizing selection, in particular, the proportion of rare and common variants present in the population. To assess enrichment/depletion of variants at specific MAF thresholds at a specific time point under stabilizing selection compared to neutrality, we constructed a 2×2 contingency table with the following groups; number of variants within MAF group, number of variants not in MAF group under neutrality and under stabilizing selection within a simulation from which an odds ratio was calculated. Application across replicates produced a distribution of odds ratios, which we log transformed and formally tested against a null hypothesis of a mean equal to zero using a standard t-test.

Next, to further investigate the contribution of rare variants to the genetic architecture across the trait continuum, we split our population into a hundred quantiles from low to high trait values. Each quantile consists of 100 individuals. For each quantile, we calculate the number of rare mutations and total number of mutations carried by individuals, both under neutrality and under selection. As described in the above paragraph, we then calculate the odds ratio within a simulation. To test if rare variants are significantly enriched in a quantile, we apply a t-test for the mean of the log-transformed odds ratio equal to zero using 100 independent simulations. Rare variants were stratified by allele frequency (MAF < 1% and MAF <= 0.05%) and effect size (all variants or top 10% with largest absolute effect size). Analyses were conducted for each group of rare variants.

### Measuring the impact of stabilising selection on PRS

To investigate how changes in the genetic architecture of the trait impacts PRS, we constructed a statistic ΔPRS that measures the differences between expected PRS and observed PRS across the trait distribution (see Fig. S2). The PRS was calculated as the sum of common variants (MAF > 1% PRS). Since we assumed heritability of 1 the phenotype is simply the sum of all variants. Both distributions were standardized by centering and scaling to mean zero and standard deviation of one. For each of the two distributions, the average was calculated per quantile (N=100 individuals per quantile). We regressed the average PRS on the average phenotype across quantiles 10-90. The expected average PRS for quantile 1-10 and 91-100, was defined as;

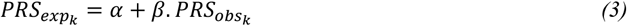

where *k* is the quantile, *PRS*_*obs*_ is the average PRS of quantile *k, α* is the intercept and *β* is the coefficient of the regression based on quantiles 10-90. ΔPRS was defined as |PRS_exp_| - |PRS_obs_| for each quantile at each time point within a simulation.

### Rescue experiments across classes of rare variants

To measure the contribution of rare variants to the tails, we recalculated PRS with the inclusion of rare variants (i.e. PRS_+rare_), which we defined across three MAF thresholds and two effect size thresholds. MAF thresholds for inclusion were: i) singletons only, ii) MAF < 0.05% (i.e. count < 11), and iii) all rare variants with MAF < 1%). Effect size thresholds were: i) any effect size and ii) top 10% based on absolute effect size. After recalculation of PRS with specific rare variants included, we also recalculated the expected average PRS in quantile 1-10 and 91-100, and subsequently also ΔPRS_+rare_. This allowed us to determine the proportion explained (*P*) in ΔPRS by each group of rare variants at each quantile (*k*) across the distribution as follows;

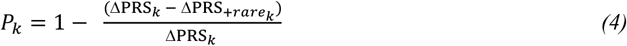

To further understand how rare variants contribute to the genetic architecture of the trait, we determined the number of individuals at the tails that carry such variants, the number of variants, and their average effect size.

## Data and resource availability

All data are produced by running the scripts publicly available from the Github: https://github.com/anilpsori/tails_simulation.

## Author’s contributions

APSO and PFO conceived and designed the study with input from CJH. CGD established the project’s first simulation pipeline in SLiM, which APSO built on and extended. APSO performed data analyses with input from CJH and PFO. APSO, CJH, and PFO performed primary interpretation of results. APSO and PFO wrote the manuscript with input from CGD and CJH, and all authors reviewed and approved the final version.

## Acknowledgements

This work was supported by NIH grants R01MH122866 and R01HG012773 (to PFO), by a Talent Grant Fellowship awarded to APSO by Amsterdam Brain and Cognition, University of Amsterdam, the Netherlands, and through the computational resources and staff expertise provided by the Data Ark and Scientific Computing teams at the Icahn School of Medicine at Mount Sinai. We are grateful for helpful discussions about this work with: Ben Haller and the SLiM team, Alanna Cote, Conrad Iyegbe, Judit García-González, Lathan Liou, Laura Sloofman, Tade Souaiaia, and Hei Man Wu.

---

**Figure S1.**
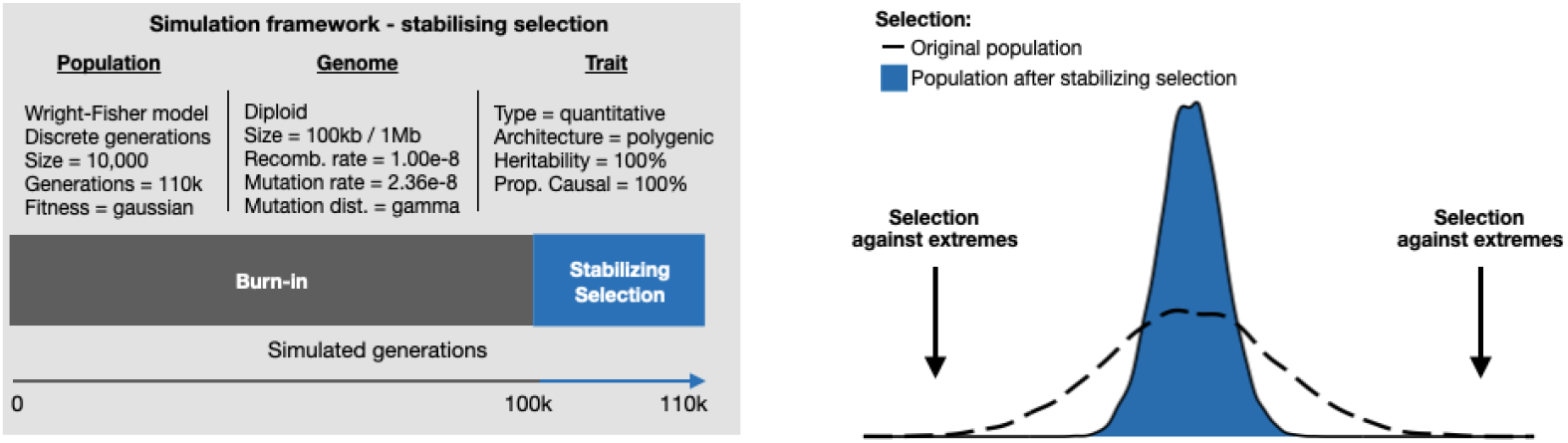
Schematic overview of WF model of stabilising selection. Shown (on the left) are key parameters that the define the population, the genome, and the trait that we have simulated. Stabilising selection, which favors the average trait value in the population and selects against individuals with trait values away from the mean (figure on the right), was introduced for a period of 1N (10,000) generations (in blue).

**Figure S2.**
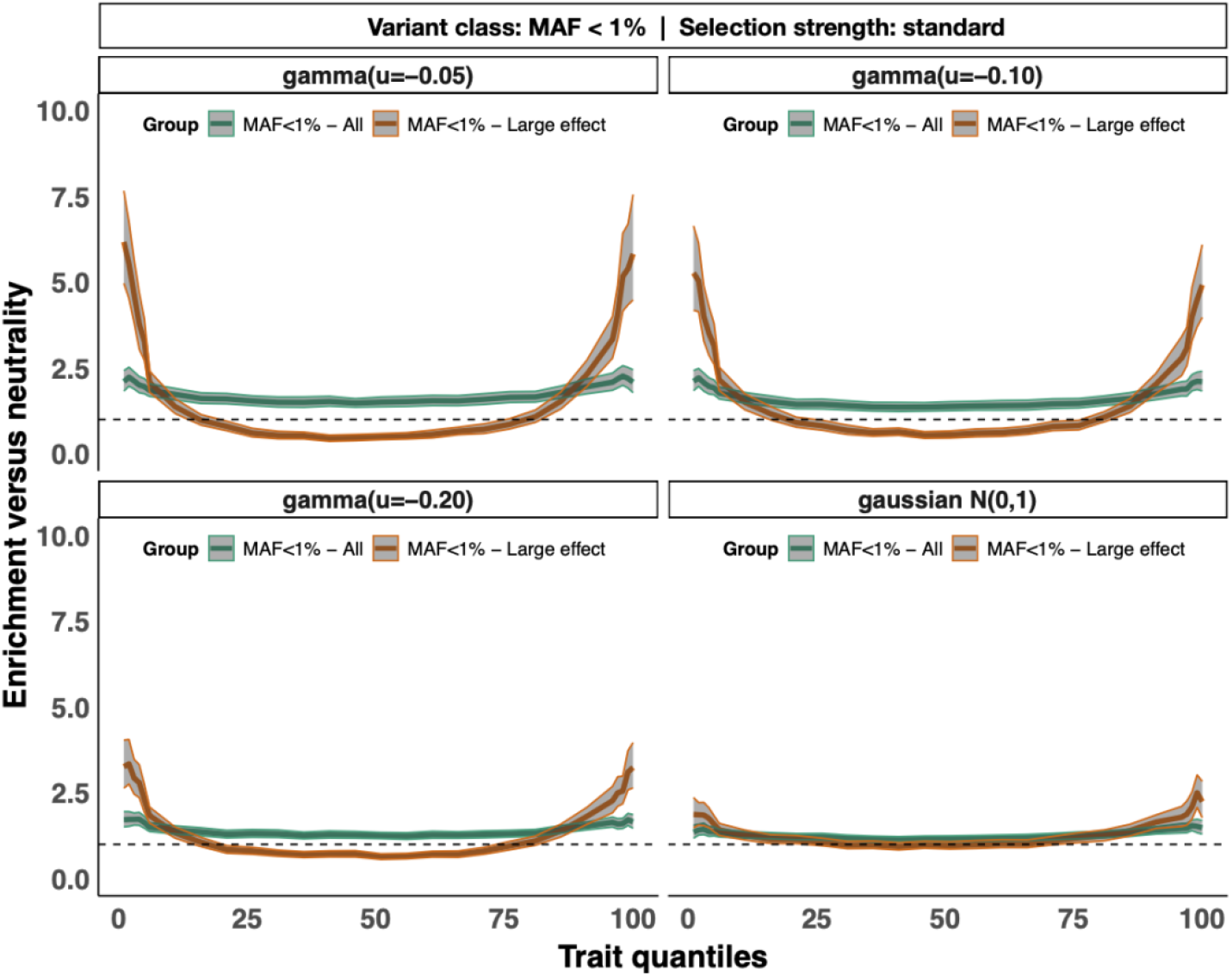
Enrichment of rare variants in the trait tails after selection stratified by different gamma and gaussian mutation generating distributions. Shown is the enrichment of rare variants (MAF < 1%) after (10k generations of) stabilising selection across quantiles of the phenotype distribution for four different mutation generation distributions; gamma with mean = −0.05 (upper left), gamma with mean = −0.10 (upper right), gamma with mean = −0.20 (bottom left), and gaussian with N(0,1) (bottom right). All three gamma distribution had a shape parameter of 0.186. Enrichment (Y-axis) is calculated, for each trait quantile, as an odds ratio defined by the ratio of rare:total alleles carried by individuals after selection divided by that ratio under neutrality (i.e. after burn-in at generation 100k). For each panel, the enrichment is shown across the trait distribution with corresponding 95% confidence intervals. In green are results for all variants with MAF < 1%, while orange shows results for variants with MAF < 1% that also have a large effect (defined as alleles with effects in the top decile of absolute effect size). The dashed horizontal lines represent an odds ratio of one (i.e. no difference between neutrality and selection).

**Figure S3.**
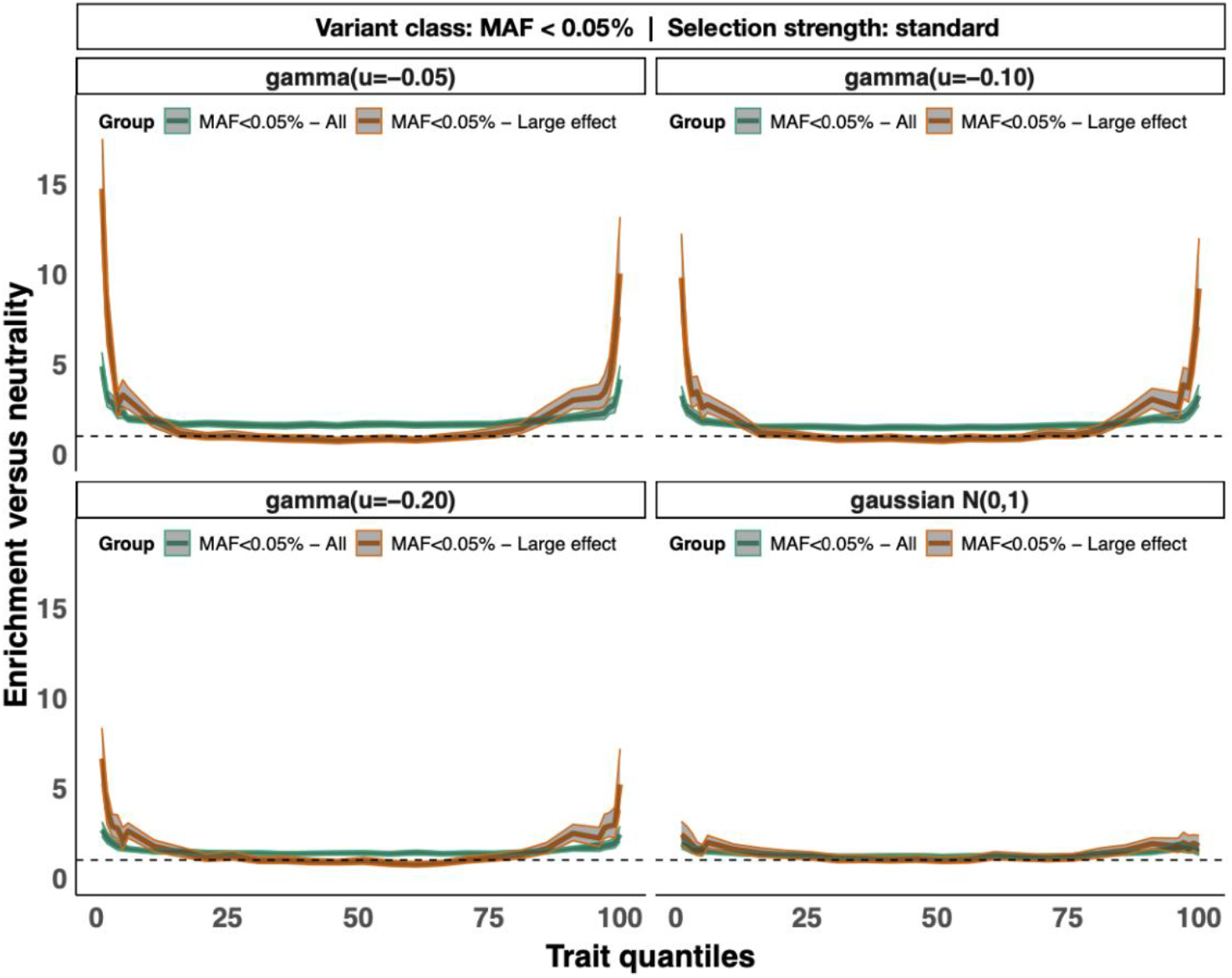
Enrichment of extremely rare variants in the trait tails after selection stratified by different gamma and gaussian mutation generating distributions. Shown is the enrichment of extremely rare variants (MAF < 0.05%) after (10k generations of) stabilising selection across quantiles of the phenotype distribution for four different mutation generation distributions; gamma with mean = −0.05 (upper left), gamma with mean =−0.10 (upper right), gamma with mean = −0.20 (bottom left), and gaussian with N(0,1) (bottom right). All three gamma distribution had a shape parameter of 0.186. Enrichment (Y-axis) is calculated, for each trait quantile, as an odds ratio defined by the ratio of rare:total alleles carried by individuals after selection divided by that ratio under neutrality (i.e. after burn-in at generation 100k). For each panel, the enrichment is shown across the trait distribution with corresponding 95% confidence intervals. In green are results for all variants with MAF < 0.05%, while orange shows results for variants with MAF < 0.05% that also have a large effect (defined as alleles with effects in the top decile of absolute effect size). The dashed horizontal lines represent an odds ratio of one (i.e. no difference between neutrality and selection).

**Figure S4.**
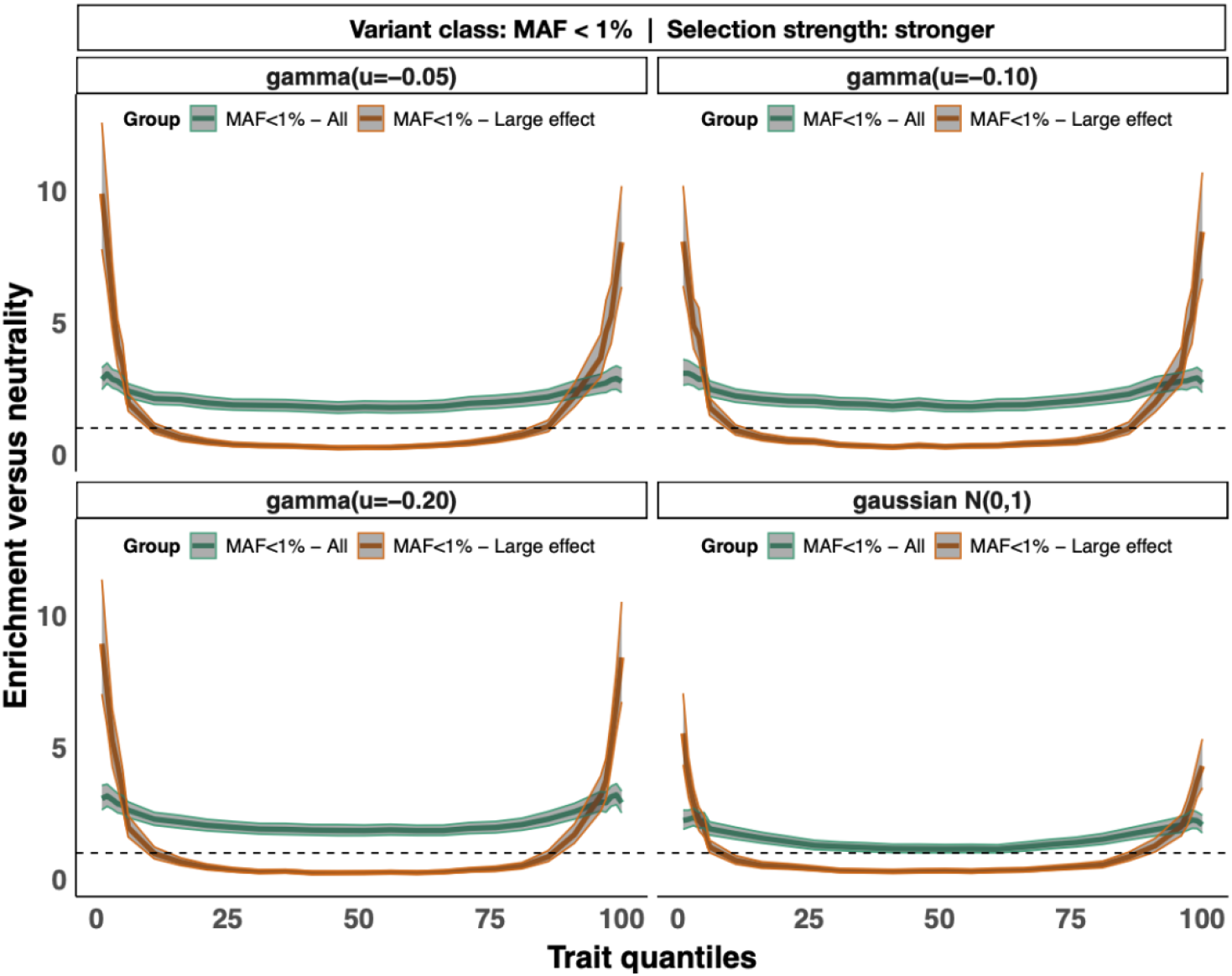
Enrichment of rare variants in the trait tails after stronger selection stratified by different gamma and gaussian mutation generating distributions. Shown is the enrichment of rare variants (MAF < 1%) after (10k generations of) stronger stabilising selection across quantiles of the phenotype distribution for four different mutation generation distributions; gamma with mean = −0.05 (upper left), gamma with mean = −0.10 (upper right), gamma with mean = −0.20 (bottom left), and gaussian with N(0,1) (bottom right). All three gamma distribution had a shape parameter of 0.186. Enrichment (Y-axis) is calculated, for each trait quantile, as an odds ratio defined by the ratio of rare:total alleles carried by individuals after stronger selection divided by that ratio under neutrality (i.e. after burn-in at generation 100k). For each panel, the enrichment is shown across the trait distribution with corresponding 95% confidence intervals. In green are results for all variants with MAF < 1%, while orange shows results for variants with MAF < 1% that also have a large effect (defined as alleles with effects in the top decile of absolute effect size). The dashed horizontal lines represent an odds ratio of one (i.e. no difference between neutrality and selection).

**Figure S5.**
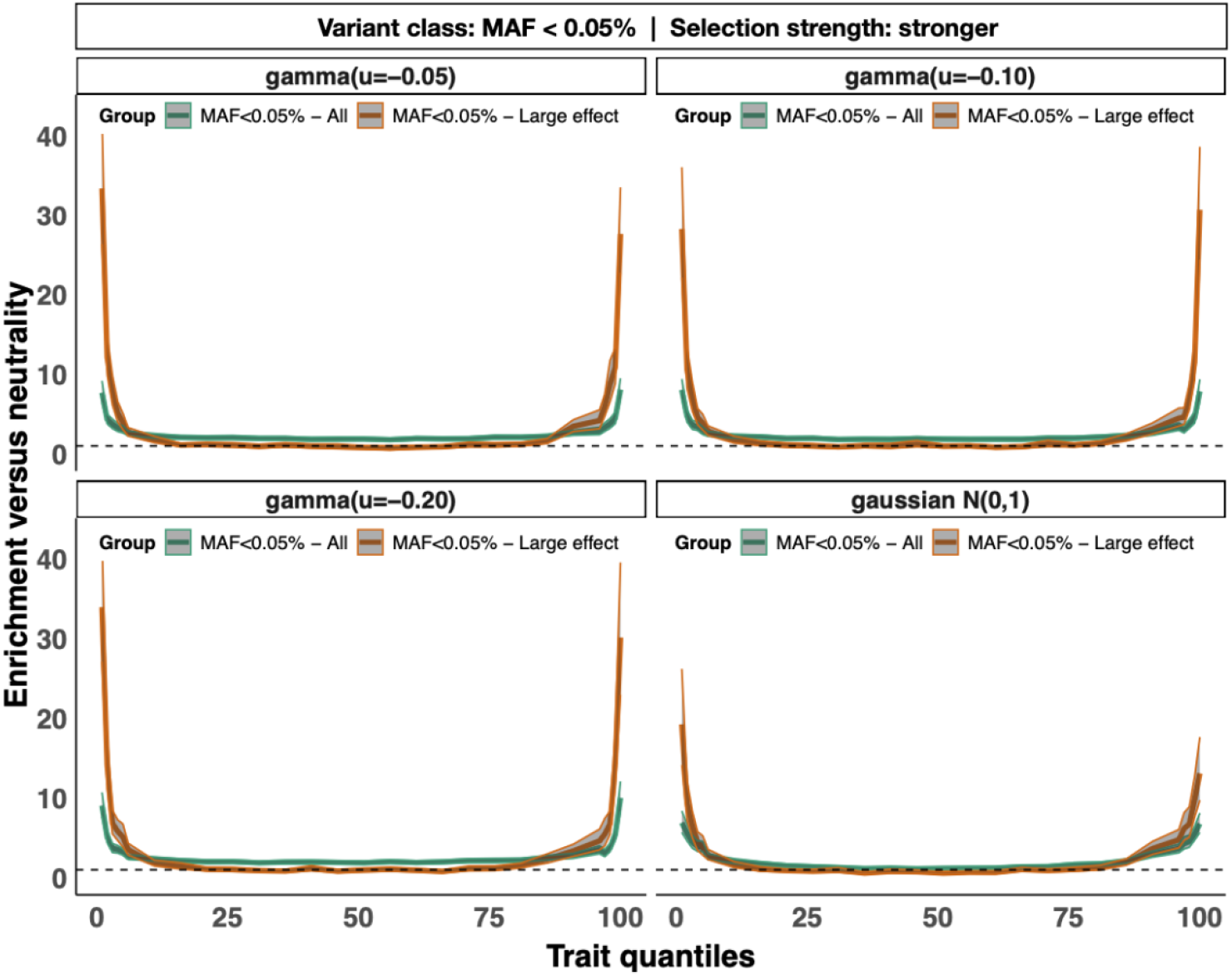
Enrichment of extremely rare variants in the trait tails after stronger selection stratified by different gamma and gaussian mutation generating distributions. Shown is the enrichment of extremely rare variants (MAF < 0.05%) after (10k generations of) stronger stabilising selection across quantiles of the phenotype distribution for four different mutation generation distributions; gamma with mean = −0.05 (upper left), gamma with mean = −0.10 (upper right), gamma with mean = −0.20 (bottom left), and gaussian with N(0,1) (bottom right). All three-gamma distribution had a shape parameter of 0.186. Enrichment (Y-axis) is calculated, for each trait quantile, as an odds ratio defined by the ratio of rare:total alleles carried by individuals after stronger selection divided by that ratio under neutrality (i.e. after burn-in at generation 100k). For each panel, the enrichment is shown across the trait distribution with corresponding 95% confidence intervals. In green are results for all variants with MAF < 0.05%, while orange shows results for variants with MAF < 0.05% that also have a large effect (defined as alleles with effects in the top decile of absolute effect size). The dashed horizontal lines represent an odds ratio of one (i.e. no difference between neutrality and selection).

**Figure S6.**
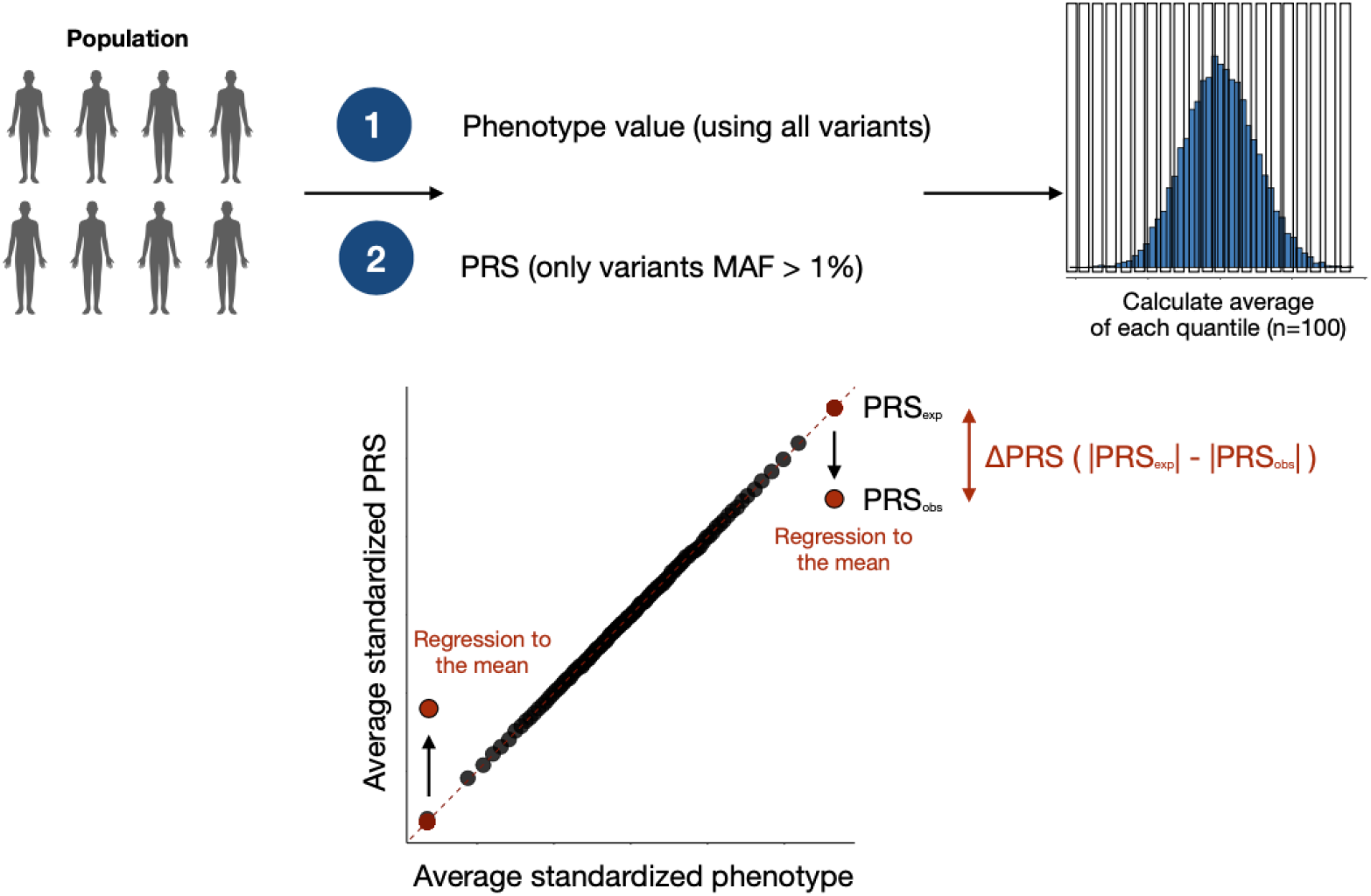
Schematic explanation of ΔPRS calculation to measure impact of rare variants at the tails. The top part of the figure shows how the distributions of the phenotype and PRS in the population are divided into 100 quantiles, for which the mean is subsequently calculated. To determine the degree of regression to the mean at the tails (i.e. ΔPRS), mean observed PRS is regressed on the mean phenotype across quantiles 11-90, for which the intercept and regression coefficients were then used to determine the mean expected PRS for each quantile. ΔPRS was calculated by subtracting the absolute expected ΔPRS (|PRS_exp_|) from the absolute observed mean observed PRS (|PRS_obs_|) for each quantile, as shown in a dummy example for a single simulation in the bottom part of the figure. If ΔPRS is close to zero, there is little to no regression to the mean effect, which indicates that rare variants carried by individuals in the tails have a similar impact away from the tails, closer to the mean of the distribution. However, if ΔPRS is significantly greater than zero, than there is a significant regression to the mean effect, indicating that rare variants carried by individuals in the tails differentially impact trait genetic architecture in the tails.

**Figure S7.**
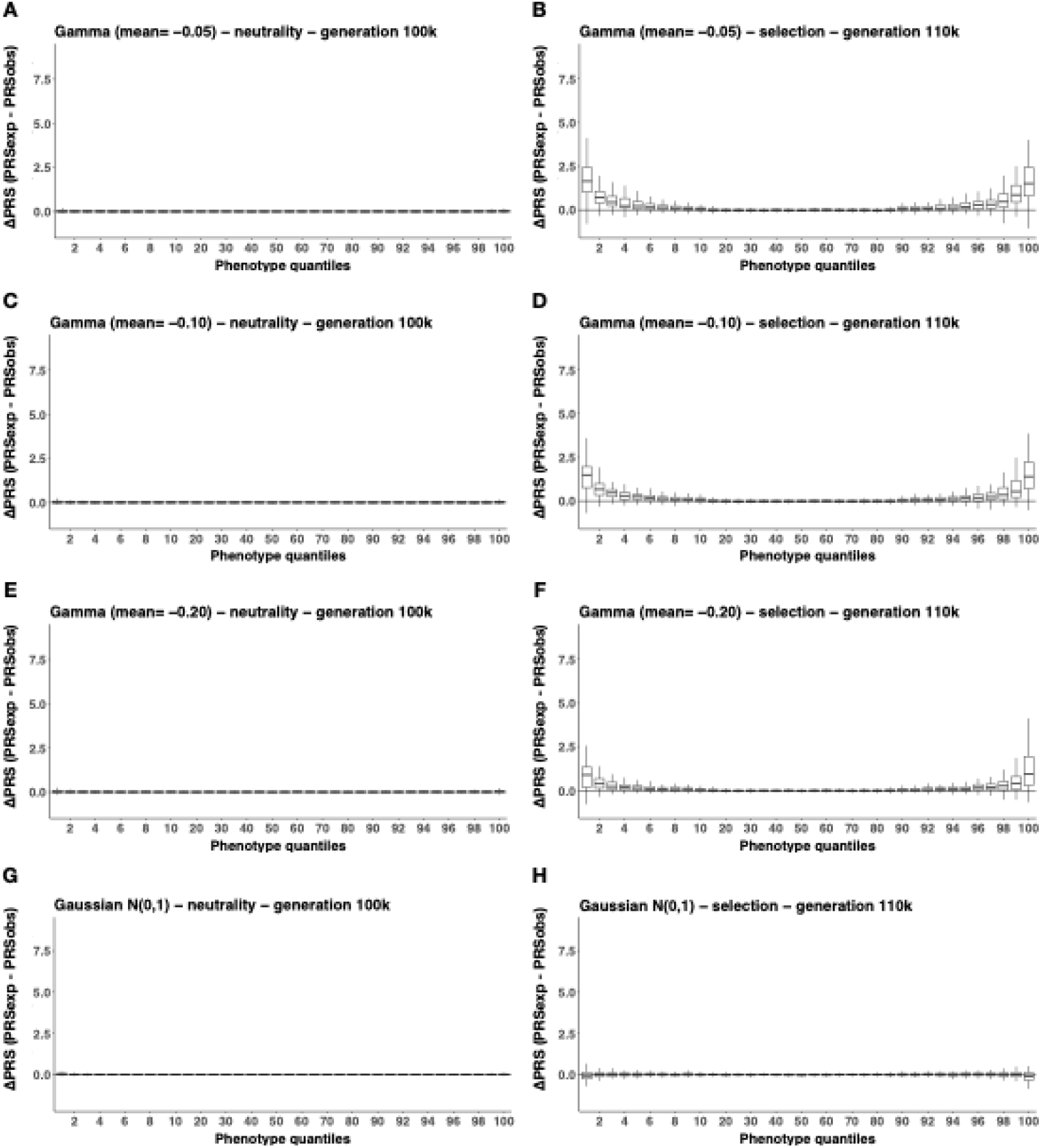
Variation in ΔPRS across the trait continuum under neutrality and stabilising selection stratified by gamma and gaussian mutation generating distributions. The average ΔPRS (see Methods) of individuals in each percentile of the trait distribution (x-axis) for 100 simulation replicates summarised as box plots. ΔPRS (y-axis) is shown under neutrality and under stabilising selection for four different mutation generating distributions; gamma with mean=−0.05 (A-B), gamma with mean=−0.10 (C-D), gamma with mean=−0.20 (E-F), and gaussian as N(0,1) (G-F). All gamma’s had a shape parameter of 0.186. ΔPRS is calculated is |PRSexp| - |PRSobs|.

**Figure S8.**
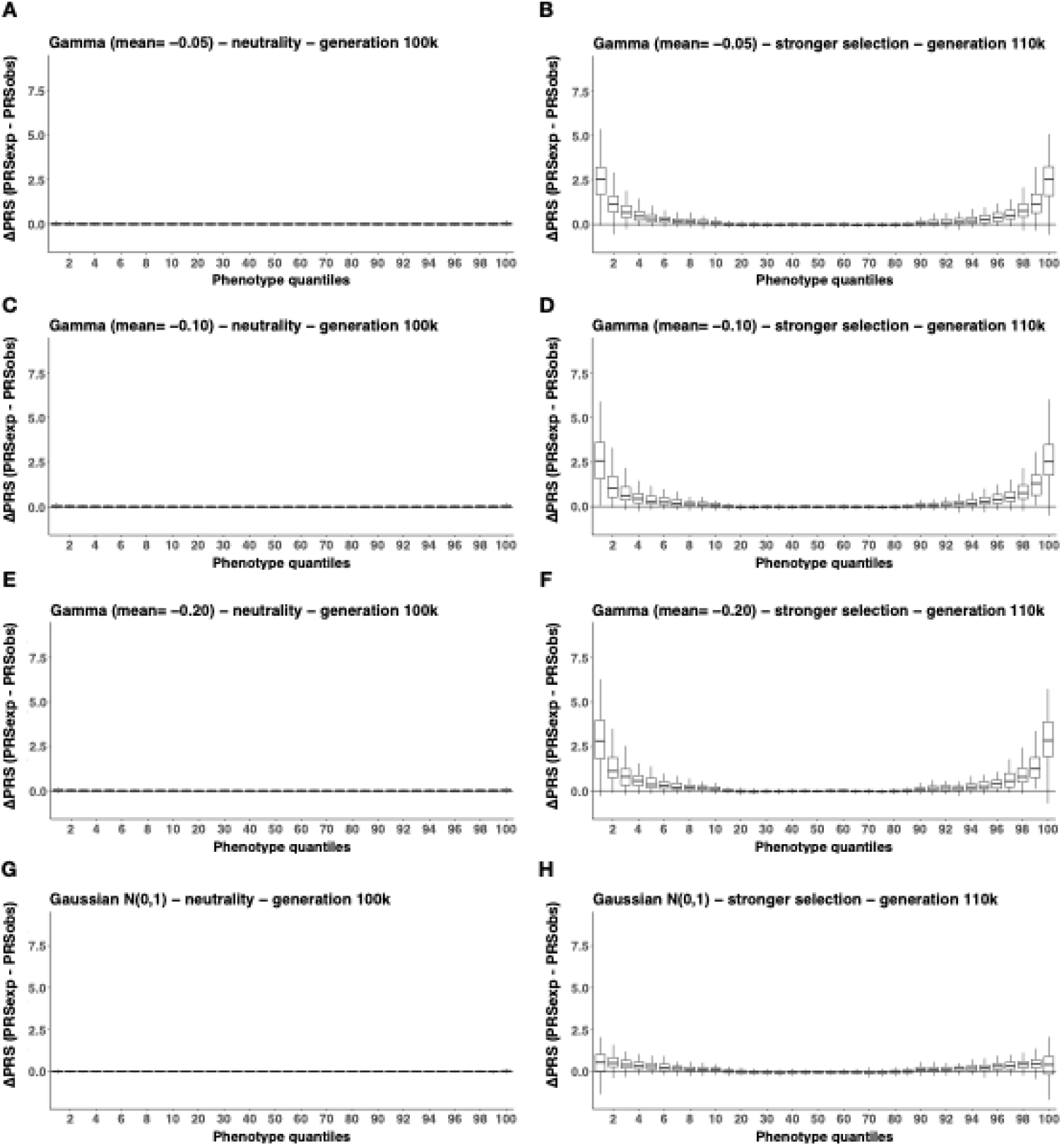
Variation in ΔPRS across the trait continuum under neutrality and stronger stabilising selection across gamma and gaussian mutation generating distributions. The average ΔPRS (see Methods) of individuals in each percentile of the trait distribution (x-axis) for 100 simulation replicates summarised as box plots. ΔPRS (y-axis) is shown under neutrality and under stronger stabilising selection for four different mutation generating distributions; gamma with mean=−0.05 (A-B), gamma with mean=−0.10 (C-D), gamma with mean=−0.20 (E-F), and gaussian as N(0,1) (G-F). All gamma’s had a shape parameter of 0.186. ΔPRS is calculated is |PRSexp| - |PRSobs|.

